# The ankyrin repeat protein RARP-1 is a periplasmic factor that supports *Rickettsia parkeri* growth and host cell invasion

**DOI:** 10.1101/2022.02.23.481736

**Authors:** Allen G. Sanderlin, Ruth E. Hanna, Rebecca L. Lamason

## Abstract

*Rickettsia* spp. are obligate intracellular bacterial pathogens that have evolved a variety of strategies to exploit their host cell niche. However, the bacterial factors that contribute to this intracellular lifestyle are poorly understood. Here, we show that the conserved ankyrin repeat protein RARP-1 supports *Rickettsia parkeri* infection. Specifically, RARP-1 promotes efficient host cell entry and growth within the host cytoplasm, but it is not necessary for cell-to-cell spread or evasion of host autophagy. We further demonstrate that RARP-1 is not secreted into the host cytoplasm by *R. parkeri*. Instead, RARP-1 resides in the periplasm, and we identify several binding partners that are predicted to work in concert with RARP-1 during infection. Altogether, our data reveal that RARP-1 plays a critical role in the rickettsial life cycle.

**Importance:** *Rickettsia* spp. are obligate intracellular bacterial pathogens that pose a growing threat to human health. Nevertheless, their strict reliance on a host cell niche has hindered investigation of the molecular mechanisms driving rickettsial infection. This study yields much needed insight into the *Rickettsia* ankyrin repeat protein RARP-1, which is conserved across the genus but has not yet been functionally characterized. Earlier work had suggested that RARP-1 is secreted into the host cytoplasm. However, the results from this work demonstrate that *R. parkeri* RARP-1 resides in the periplasm and is important both for invasion of host cells and for growth in the host cell cytoplasm. These results reveal RARP-1 as a novel regulator of the rickettsial life cycle.

## Introduction

Intracellular bacterial pathogens face considerable challenges and opportunities when invading and occupying their host cell niche. The host cell membrane physically occludes entry and the endolysosomal pathway imperils invading microbes. Moreover, host cell defenses like autophagy create a hostile environment for internalized bacteria. If a bacterium successfully navigates these obstacles, however, it can conceal itself from humoral immunity, commandeer host metabolites, and exploit host cell biology to support infection. Not surprisingly, the host cell niche has provided fertile ground for the evolution of diverse lifestyles across many well-studied bacterial pathogens such as *Shigella*, *Listeria*, *Salmonella*, and *Legionella* (1, 2). The prospect of uncovering unique infection strategies invites a thorough investigation of these adaptations in more enigmatic pathogens.

Members of the genus *Rickettsia* include emerging global health threats that can cause mild to severe diseases such as typhus and Rocky Mountain spotted fever (3). These Gram-negative bacterial pathogens are transmitted from arthropod vectors to vertebrate hosts where they primarily target the vascular endothelium. As obligate intracellular pathogens, *Rickettsia* spp. define the extreme end of adaptation to intracellular life and are completely dependent on their hosts for survival (4). Consequently, they have evolved a complex life cycle to invade, grow, and disseminate across host tissues.

As the first step of their life cycle, *Rickettsia* spp. adhere to and invade host cells by inducing phagocytosis (5–7). Once inside, these bacteria rapidly escape the phagocytic vacuole to access the host cytoplasm (8, 9). To establish a hospitable niche for proliferation, *Rickettsia* spp. scavenge host nutrients, modulate apoptosis, and thwart antimicrobial autophagy (10–13). Successful colonization of the host cytoplasm allows *Rickettsia* spp. to spread to neighboring cells. Members of the spotted fever group (SFG) *Rickettsia* hijack the host actin cytoskeleton, forming tails that propel the bacteria around the cytoplasm, and then protrude through cell-cell junctions to repeat the infection cycle (14, 15).

Recent work using the model SFG member *Rickettsia parkeri* has highlighted a short list of surface-exposed proteins and secreted effectors that manipulate host cell processes during infection (4). For example, the surface protein Sca2 nucleates actin at the bacterial pole and promotes motility by mimicking host formins (14). Sca4, a secreted effector, interacts with host vinculin to reduce intercellular tension and facilitate protrusion engulfment (15). Additionally, methylation of outer membrane proteins like OmpB protects *R. parkeri* from ubiquitylation and autophagy (13, 16). Despite these advances, our knowledge of the factors that govern the multi-step rickettsial life cycle is still limited. Indeed, *Rickettsia* spp. genomes are replete with hypothetical proteins that are conserved even among less virulent members of the genus (17), but a paucity of genetic tools has stunted investigation of these proteins. Such factors could support infection directly, by targeting host processes, or indirectly, by controlling the bacterial mediators at the host-pathogen interface. Thus, it is critical to reveal how these uncharacterized proteins contribute to infection.

In a recent transposon mutagenesis screen in *R. parkeri* (18), we identified over 100 mutants that exhibited defects in infection. Although several hits from this screen have been functionally characterized (13–16), many play unknown roles during infection. One such unexplored hit is the *Rickettsia* ankyrin repeat protein 1 (RARP-1), which is conserved across the genus and predicted to be secreted into the host cytoplasm (19). To better understand the factors that influence the rickettsial life cycle, we investigated the function of RARP-1 during *R. parkeri* infection. We demonstrated that RARP-1 promotes both efficient host cell invasion and growth in the host cytoplasm, but it is otherwise dispensable for cell-to-cell spread and avoidance of host autophagy. Although prior work indicated that RARP-1 is secreted into the host cytoplasm (19), we found instead that it localizes to the *R. parkeri* periplasm. Furthermore, we showed that RARP-1 interacts with a variety of factors that are predicted to support bacterial fitness. Our results suggest that RARP-1 is a *Rickettsia*-specific tool that promotes the obligate intracellular life cycle.

## Results

### Transposon mutagenesis of *rarp-1* impairs *R. parkeri* infection

In a previous *mariner*-based transposon mutagenesis screen (18), we identified a number of *R. parkeri* mutants that displayed abnormal plaque sizes after infection of Vero host cell monolayers. We hypothesized that the plaque phenotypes for these mutants were due to defects in growth, cell-to-cell spread, or other steps of the rickettsial life cycle. Two such small plaque (Sp) mutants contained a transposon (Tn) insertion within the *rarp-1* gene, giving a predicted truncation of RARP-1 at residues 305 (Sp116) and 480 (Sp64) (Figure 1A). RARP-1 is a 573 amino acid protein conserved across the *Rickettsia* genus, but the lack of loss-of-function mutants has thus far prevented characterization of RARP-1 function. Due to the upstream position of its Tn insertion within the *rarp-1* CDS, we focused on Sp116 (herein referred to as *rarp-1*::Tn) for all subsequent studies and confirmed that it formed smaller plaques than GFP-expressing wild-type bacteria (WT, Figure 1B). We generated polyclonal antibodies against a RARP-1 peptide upstream of the Tn insertion site to assess RARP-1 expression in the mutant. As expected, the *rarp-1*::Tn mutant did not express the full-length protein by immunoblotting (Figure 1C). Furthermore, we were unable to detect an obvious band consistent with the expected 30 kDa product resulting from Tn insertion. Altogether, these results suggest that the loss of RARP-1 expression in the *rarp-1*::Tn mutant leads to a small plaque phenotype.

**Figure 1.**
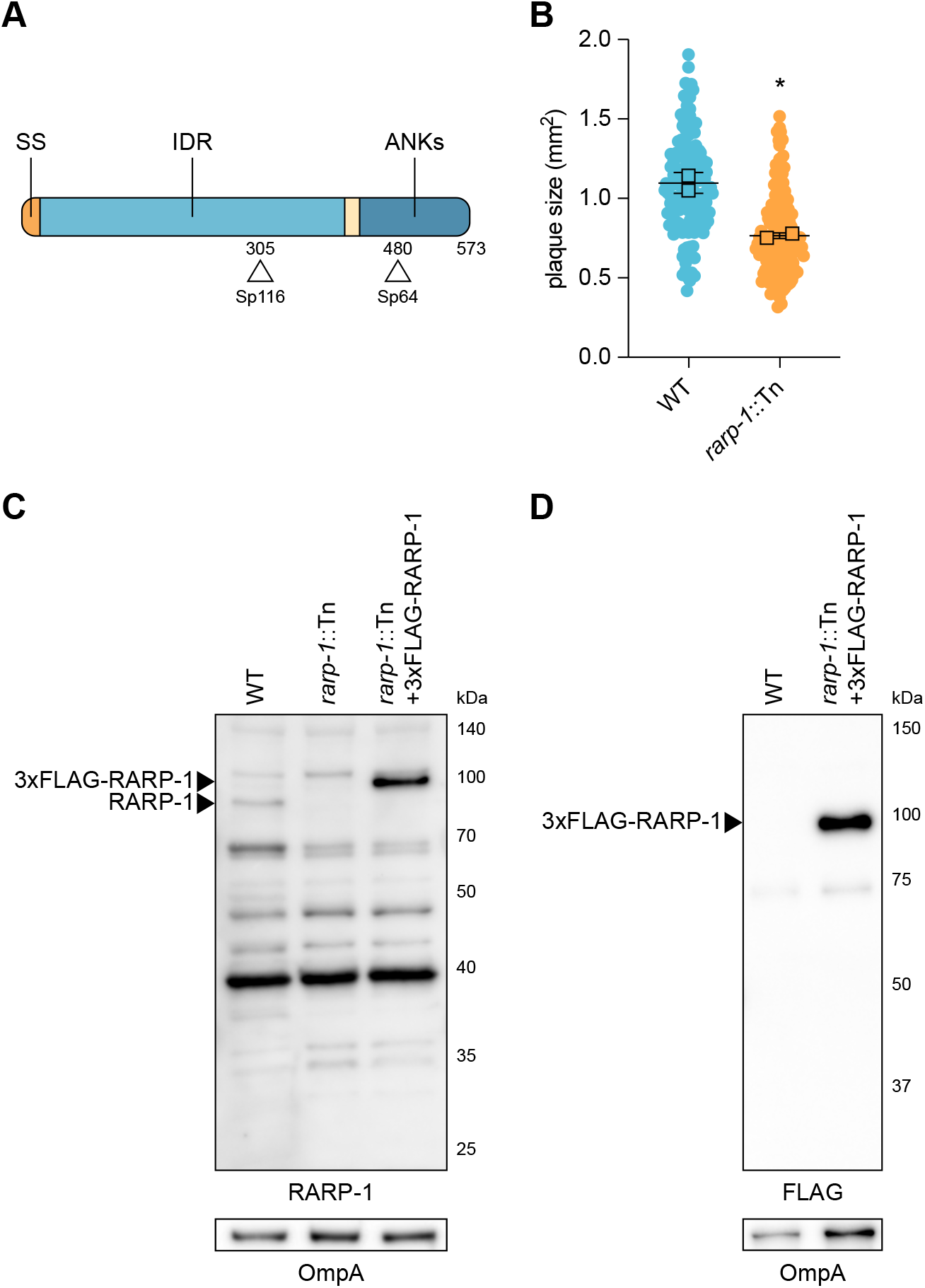
Transposon mutagenesis of *rarp-1* impairs *R. parkeri* infection. (A) *R. parkeri* RARP-1 contains an N-terminal Sec secretion signal (SS, orange), a central intrinsically disordered region (IDR, light blue), and C-terminal ankyrin repeats (ANKs, dark blue). Tn insertions at residues 305 (Sp116) and 480 (Sp64) are indicated (arrowheads). (B) Plaque areas in infected Vero cell monolayers. Means from two independent experiments (squares) are superimposed over the raw data (circles) and were used to calculate the mean ± SD and p-value (unpaired two-tailed *t* test, *p < 0.05 relative to WT). (C) Western blot for RARP-1 using purified *R. parkeri* strains. 3xFLAG-tagged and endogenous RARP-1 are indicated (arrowheads). OmpA, loading control. (D) Western blot for FLAG using purified *R. parkeri* strains. 3xFLAG-tagged RARP-1 is indicated (arrowhead). OmpA, loading control.

### RARP-1 supports bacterial growth and is dispensable for cell-to-cell spread

The small plaques formed by the *rarp-1*::Tn mutant could be the result of defects in one or more steps of the rickettsial life cycle, and determining when RARP-1 acts during infection would support characterization of its function. We first performed infectious focus assays in A549 host cell monolayers to assess the growth and cell-to-cell spread of the *rarp-1*::Tn mutant on a shorter timescale than is required for plaque formation (28 h versus 5 d post-infection). A549 cells support all known aspects of the *R. parkeri* life cycle, and the use of gentamicin prevents asynchronous invasion events (15). Consistent with the small plaque phenotype, the *rarp-1*::Tn mutant generated smaller foci than WT bacteria (Figure 2A). To confirm that this phenotype was due specifically to the disruption of *rarp-1*, we complemented the *rarp-1*::Tn mutant with a plasmid expressing 3xFLAG-tagged RARP-1 (*rarp-1*::Tn + 3xFLAG-RARP-1, Supplementary Figure 1A). Since *rarp-1* is predicted to be part of an operon (19), we selected a 247 bp region immediately upstream of the first gene in the operon (encoding the outer membrane channel TolC) as a putative promoter to drive *rarp-1* expression. This construct was sufficient for expression of epitope-tagged RARP-1 in the *rarp-1*::Tn mutant (Figures 1C and D). Importantly, the complement strain exhibited infectious focus sizes comparable to WT (Figure 2A), indicating that the putative promoter and epitope-tagged RARP-1 are functionally relevant. Thus, RARP-1 specifically supports the size of *R. parkeri* infectious foci.

**Figure 2.**
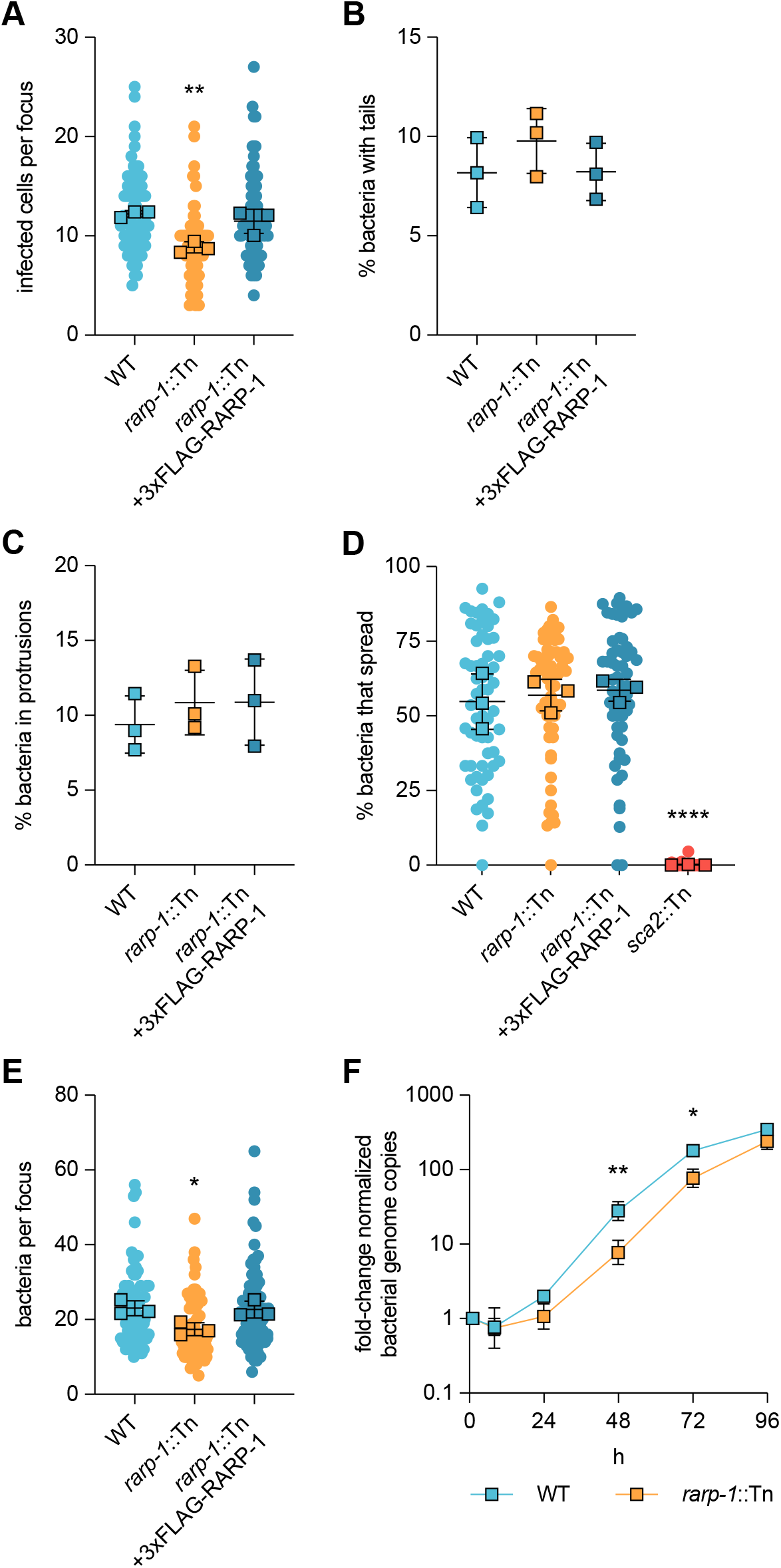
RARP-1 supports bacterial growth and is dispensable for cell-to-cell spread. (A) Infected cells per focus during infection of A549 cells. The means from three independent experiments (squares) are superimposed over the raw data (circles) and were used to calculate the mean ± SD and p-value (one-way ANOVA with post-hoc Dunnett’s test, **p < 0.01 relative to WT). (B) Percentage of bacteria with actin tails during infection of A549 cells. (C) Percentage of bacteria within a protrusion during infection of A549 cells. In (B) and (C), the percentages were determined from three independent experiments (≥ 380 bacteria were counted for each infection) and were used to calculate the mean ± SD and p-value (one-way ANOVA with post-hoc Dunnett’s test, n.s. relative to WT). (D) Percentage of bacteria per focus that spread from infected donor cells to uninfected recipient cells by mixed-cell assay in A549 cells. The means from three independent experiments (squares) are superimposed over the raw data (circles) and were used to calculate the mean ± SD and p-value (one-way ANOVA with post-hoc Dunnett’s test, ****p < 0.0001 relative to WT). The *sca2*::Tn mutant was used as a positive control. (E) Bacteria per focus during infection of A549 cells. The means from three independent experiments (squares) are superimposed over the raw data (circles) and were used to calculate the mean ± SD and p-value (one-way ANOVA with post-hoc Dunnett’s test, *p < 0.05). These data correspond to the same set of infectious focus assays displayed in (A). (F) Growth curves as measured by *R. parkeri* (17 kDa surface antigen) genome equivalents per Vero host cell (*GAPDH*) genome equivalent normalized to 1 h post-infection. The mean ± SD for triplicate samples from a representative experiment were compared at each timepoint after log_2_ transformation (unpaired two-tailed *t* test, *p < 0.05 and **p < 0.01 relative to WT).

A reduction in infectious focus size could be caused by defects in cell-to-cell spread. For example, Tn mutagenesis of *sca2* and *sca4* specifically disrupts spread by limiting actin tail formation and protrusion resolution, respectively, leading to smaller infectious foci (14, 15). Loss of RARP-1 did not alter the frequency of actin tails or protrusions (Figures 2B and C), suggesting that spread may not be regulated by RARP-1. As an orthogonal approach, we also evaluated the efficiency of spread by performing a mixed-cell infectious focus assay (15). In this assay, donor host cells stably expressing a cytoplasmic marker are infected, mixed with unlabeled recipient host cells, and then infection of the mixed monolayer is allowed to progress. Bacteria that spread to unlabeled recipient cells can thus be distinguished from bacteria that remain in the labeled donor cell for each focus. As expected, a *sca2*::Tn mutant failed to spread from infected donor cells (Figure 2D). In contrast, the *rarp-1*::Tn mutant exhibited similar efficiency of spread from donors to recipients as compared to WT bacteria. Altogether, these results indicate that RARP-1 is dispensable for cell-to-cell spread.

Alternatively, a reduction in infectious focus size could be caused by defects in bacterial growth. When performing the infectious focus assays, we noted that the number of *rarp-1*::Tn mutant bacteria within the infectious foci was reduced compared to WT (Figure 2E). This was in contrast to Tn mutants of *sca2* and *sca4*, which do not exhibit reduced bacterial loads despite forming smaller foci (14, 15). Restoring RARP-1 expression in the complement strain rescued the bacterial load defect (Figure 2E), suggesting that RARP-1 regulates bacterial growth. To determine if the *rarp-1*:Tn mutant displayed altered growth behavior over the course of infection, we used qPCR to monitor bacterial genome equivalents during infection of Vero host cell monolayers. In agreement with the bacterial load defect observed in the infectious focus assay, the *rarp-1*::Tn mutant exhibited a growth defect compared to WT (Figure 2F). Together, our data support a role for RARP-1 during bacterial growth in multiple cell types.

### RARP-1 is dispensable for evasion of autophagy

Given the *rarp-1*::Tn mutant growth defect, we hypothesized that RARP-1 might promote bacterial growth by preventing clearance from the host cell. *R. parkeri* avoids recognition and destruction by the host cell autophagy machinery using the abundant outer membrane protein OmpB (13). Bacteria lacking OmpB are readily polyubiquitinated by the host cell and associate with LC3-positive autophagic membranes. We tested whether the *rarp-1*::Tn mutant likewise associates with LC3 during infection of A549 cells. In contrast to an *ompB*::Tn mutant, the *rarp-1*::Tn mutant failed to mobilize host LC3 (Figure 3A). Thus, loss of RARP-1 expression does not render this mutant more susceptible to autophagic clearance, indicating that RARP-1 supports growth through a different mechanism.

**Figure 3.**
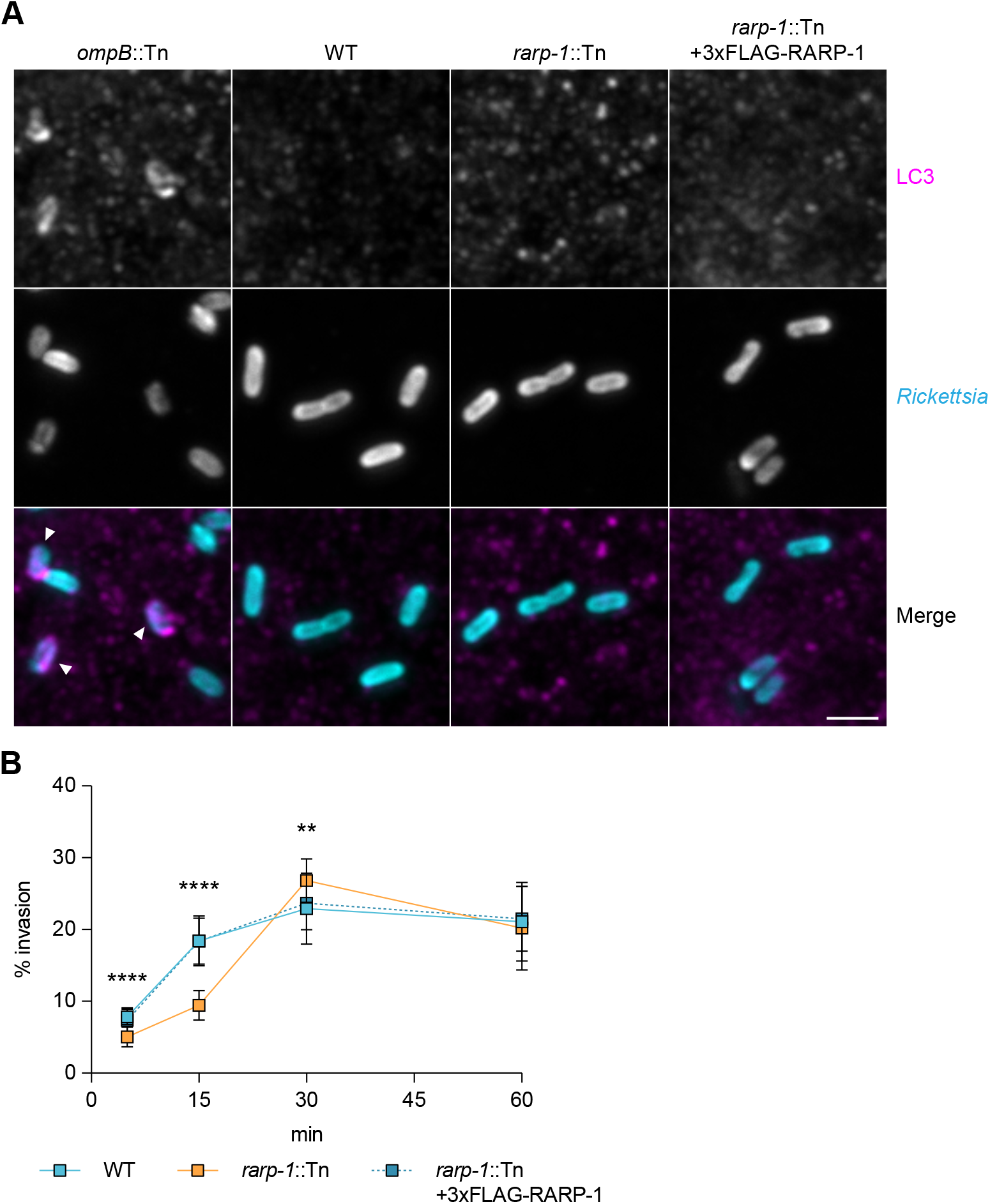
RARP-1 is dispensable for evasion of host cell autophagy and supports host cell invasion. (A) Recruitment of LC3 during infection of A549 cells. Samples were stained for LC3 (magenta) and bacteria (cyan). The *ompB*::Tn mutant was used as a positive control, and bacteria associated with LC3-positive membranes are indicated (arrowheads). Scale bar, 2 μm. (B) Efficiency of invasion into A549 cells. The means ± SD from a representative experiment (n = 20 fields of view each with ≥ 45 bacteria) were compared at each timepoint (one-way ANOVA with post-hoc Dunnett’s test, **p < 0.01 and ****p < 0.0001 relative to WT).

### RARP-1 supports host cell invasion

We next wanted to determine if RARP-1 plays other roles in the infection cycle upstream of growth inside the host cytoplasm. We tested whether the *rarp-1*::Tn mutant exhibited defects during invasion of A549 host cells using differential immunofluorescent staining (6). In this assay, bacteria are stained both before and after host cell permeabilization to distinguish external and internal bacteria, respectively. Invasion of the *rarp-1*::Tn mutant was delayed compared to WT but otherwise recovered within 30 min post-infection (Figure 3B). We observed similar invasion kinetics for WT bacteria and the complement strain, indicating that the delayed invasion of the *rarp-1*::Tn mutant is due to loss of RARP-1 expression. Thus, RARP-1 supports efficient host cell invasion. We therefore turned our investigation to the localization and binding partners of RARP-1 so that we could reveal how this factor contributes to infection.

### RARP-1 is not secreted into the host cytoplasm by *R. parkeri*

RARP-1 contains an N-terminal Sec secretion signal and several C-terminal ankyrin repeats. Ankyrin repeats are often involved in protein-protein interactions (20), and various intracellular pathogens secrete ankyrin repeat-containing proteins to target an array of host cell processes (21, 22). Previous work with the typhus group *Rickettsia* species *R. typhi* suggested that RARP-1 is delivered into host cells through a non-canonical mechanism mediated by the Sec translocon and TolC (19). We originally hypothesized that *R. parkeri* also secretes RARP-1 to target host cell functions and ultimately promote bacterial growth and invasion. To monitor secretion of RARP-1 during infection of A549 cells, we used selective lysis to separate supernatants containing the infected host cytoplasm from pellets containing intact bacteria. A protein that is secreted during infection should be detected in both the supernatant and pellet fractions by immunoblotting, as was observed for the secreted effector Sca4 (Figure 4A, middle panel). The absence of the bacterial RNA polymerase subunit RpoA in the supernatant fraction confirmed that our lysis conditions did not cause bacterial lysis and release of non-secreted bacterial proteins (Figure 4A, bottom panel). Unexpectedly, we detected 3xFLAG-RARP-1 in the bacterial pellet but not in the supernatant fraction of cells infected with the *rarp-1*::Tn + 3xFLAG-RARP-1 complement strain (Figure 4A, top panel). Similar results were observed for a 3xFLAG-RARP-1 construct containing an additional Ty1 epitope tag inserted proximal to the C-terminus (Supplementary Figure 1A), suggesting that the lack of detection was not due to proteolytic processing of the RARP-1 protein. As with the 3xFLAG-RARP-1 construct, this dual-tagged variant rescued the *rarp-1*::Tn mutant infectious focus defects (Supplementary Figures 1B and C), demonstrating the functional relevance of the tagged RARP-1 construct. Moreover, endogenous RARP-1 protein was detectable in the WT bacterial pellet but not in the supernatant fraction with our polyclonal antibody (Supplementary Figure 1D), confirming that the epitope-tagged constructs recapitulate the behavior of the endogenous protein. Together, these results suggest that RARP-1 is not secreted by *R. parkeri* into the host cytoplasm.

**Figure 4.**
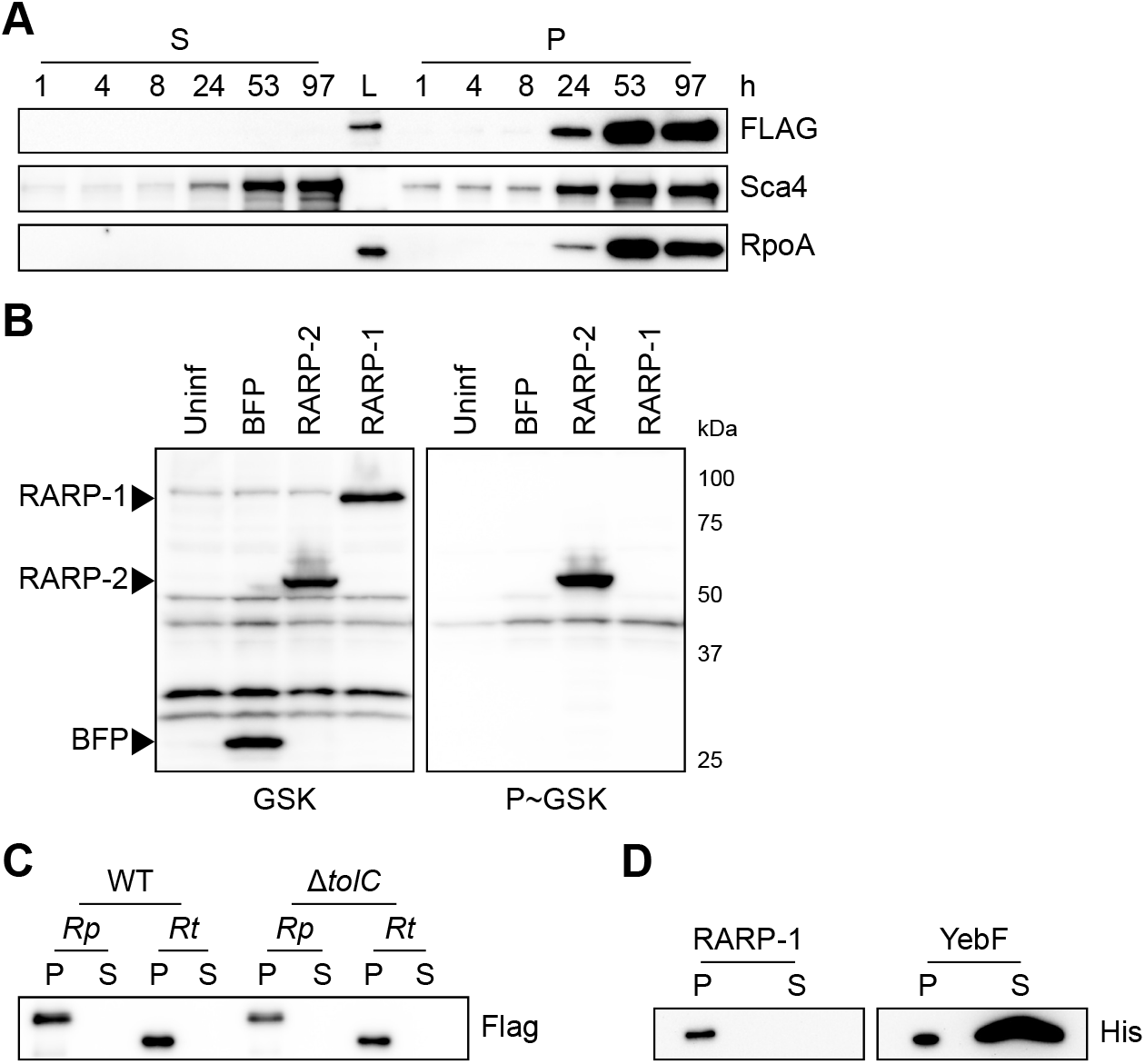
RARP-1 is not secreted. (A) Western blots for FLAG (top) and Sca4 (middle) during infection of A549 cells with *rarp-1*::Tn + 3xFLAG-RARP-1 bacteria. Infected host cells were selectively lysed at various timepoints to separate supernatants (S) containing the infected host cytoplasm from pellets (P) containing intact bacteria. RpoA (bottom) served as a control for bacterial lysis or contamination of the infected cytoplasmic fraction. L, ladder. (B) Western blot for GSK-tagged constructs during infection of Vero cells. Whole cell infected lysates were probed with antibodies against the GSK tag (left) or its phosphorylated form (P∼GSK, right) to detect exposure to the host cytoplasm. BFP (non-secreted) and RARP-2 (secreted) were used as controls. Uninf, uninfected whole cell lysate. (C) Western blot for FLAG using N-terminal FLAG-tagged *R. parkeri* (*Rp*) or *R. typhi* (*Rt*) RARP-1 expressed by WT or Δ*tolC E. coli*. (D) Western blot for His using C-terminal Myc-6xHis-tagged *R. typhi* RARP-1 or C-terminal 6xHis-tagged *E. coli* YebF expressed by WT *E. coli*. For (C) and (D), cultures were pelleted (P) and the culture supernatant (S) was filtered and precipitated to concentrate proteins released into the medium.

As an alternative strategy to evaluate RARP-1 secretion, we introduced glycogen synthase kinase (GSK)-tagged constructs into *R. parkeri*. This system has been used to assess secretion of effector proteins by *Rickettsia* spp. and other bacteria, and it does not rely on the selective lysis of infected samples (23, 24). GSK-tagged proteins become phosphorylated by host kinases upon entering the host cytoplasm, and secretion of the tagged protein can be validated by phospho-specific antibodies (25). Although GSK-tagged RARP-2, a known secreted effector (23), was phosphorylated, GSK-tagged RARP-1 and a non-secreted control (BFP) were not phosphorylated during infection (Figure 4B). These results provide further evidence that RARP-1 is not secreted into the host cytoplasm by *R. parkeri*.

### Heterologously expressed RARP-1 is not secreted by *E. coli*

We were surprised by the results above since previous work suggested that RARP-1 is delivered into host cells by *R. typhi*. Heterologous expression in *Escherichia coli* provided evidence that *R. typhi* RARP-1 is secreted in a Sec- and TolC-dependent manner (19). Following the methodology described by that work, we assessed secretion of *R. parkeri* and *R. typhi* RARP-1 by WT and Δ*tolC E. coli*. In this assay, *E. coli* cultures expressing RARP-1 are pelleted and the culture supernatant is then filtered and precipitated to concentrate proteins released into the extracellular milieu. Although *R. parkeri* RARP-1 was clearly detectable in the bacterial pellets of both strains, it was not observed in the supernatants for either strain (Figure 4C). Likewise, we were unable to detect secretion of *R. typhi* RARP-1 by either strain, in contrast to the previously described secretion pattern for this protein. To confirm that our use of an N-terminal 3xFLAG tag did not disrupt secretion by *E. coli*, we generated an *R. typhi* RARP-1 construct with a C-terminal Myc-6xHis tag, as described in the previous work. Again, we were unable to detect secretion of *R. typhi* RARP-1 (Figure 4D). To validate our ability to detect secreted proteins in the culture supernatant, we assessed secretion of 6xHis-tagged YebF, a protein known to be exported into the medium by *E. coli* (26). As expected, YebF was observed in both the bacterial pellet and culture supernatant. The lack of RARP-1 secretion by *E. coli* is consistent with our immunoblotting results for infection with *R. parkeri*, suggesting that RARP-1 is not a secreted effector.

### RARP-1 localizes to the *R. parkeri* periplasm

Given that RARP-1 is not secreted by *R. parkeri*, we next investigated where it localized during infection using differential immunofluorescent staining (15). In this assay, infected A549 host cells are first selectively permeabilized such that only the host cell contents and bacterial surface are accessible for staining. Then, the bacteria are permeabilized with lysozyme and detergent to permit immunostaining of proteins inside the bacteria. By staining with a FLAG tag-specific antibody either with or without this second permeabilization step, we can distinguish the localization of tagged proteins inside or outside the bacteria, respectively. We predicted that epitope-tagged RARP-1 expressed by the *rarp-1*::Tn + 3xFLAG-RARP-1 complement strain would be absent from the host cytoplasm but present inside permeabilized bacteria. In agreement with our immunoblotting results above, we did not detect specific FLAG staining in the host cytoplasm after infection with the complement strain, similar to results with the *rarp-1*::Tn mutant (Figure 5A). We also did not detect the protein on the bacterial surface. Instead, 3xFLAG-RARP-1 was only detectable after permeabilizing bacteria with lysozyme and detergent. Under these conditions, the 3xFLAG-RARP-1 signal surrounded the bacteria with variable localization patterns and often formed bipolar puncta (Figure 5B). Line scan analysis of permeabilized bacteria confirmed that 3xFLAG-RARP-1 localized adjacent to the bacterial cytoplasm (Figure 5C). These localization patterns, together with the presence of an N-terminal Sec secretion signal, suggest that RARP-1 is not secreted into the host cytoplasm but instead localizes to the *R. parkeri* periplasm.

**Figure 5.**
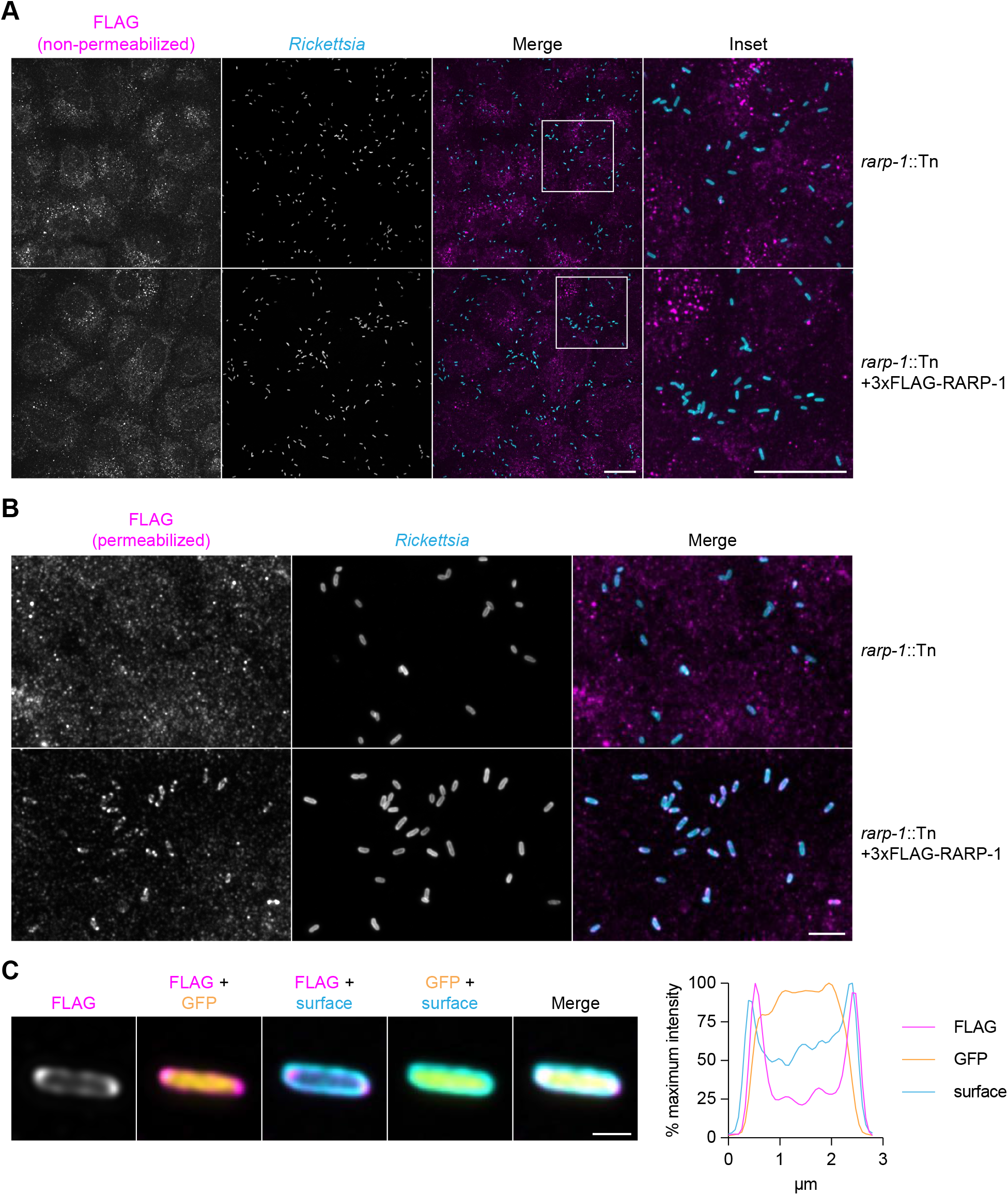
RARP-1 localizes to the *R. parkeri* periplasm. (A) Images of *rarp-1*::Tn (top) and *rarp-1*::Tn + 3xFLAG-RARP-1 (bottom) bacteria during infection of A549 cells. Samples were stained for FLAG (magenta) and the bacterial surface (cyan) without permeabilization of bacteria. Scale bars, 20 μm. (B) Images of *rarp-1*::Tn (top) and *rarp-1*::Tn + 3xFLAG-RARP-1 (bottom) bacteria during infection of A549 cells. The bacterial surface (cyan) was stained prior to permeabilization by lysozyme and detergent and subsequent staining for FLAG (magenta). Scale bar, 5 μm. (C) Subcellular localization of 3xFLAG-RARP-1 in a representative *rarp-1*::Tn + 3xFLAG-RARP-1 bacterium during infection of A549 cells. The bacterial surface (cyan) was stained prior to permeabilization by lysozyme and detergent and subsequent staining for FLAG (magenta). GFP (yellow) demarcates the bacterial cytoplasm. Scale bar, 1 μm. A pole-to-pole 0.26 μm width linescan (right) was generated for FLAG, GFP, and the bacterial surface.

### RARP-1 interacts with other bacterial factors that access the periplasm

Based on the 3xFLAG-RARP-1 localization pattern, we hypothesized that RARP-1 might interact with other factors in the *R. parkeri* periplasm to support growth and host cell invasion. To test this hypothesis, we isolated *rarp-1*::Tn + 3xFLAG-Ty1-RARP-1 bacteria and treated them with lysozyme-containing lysis buffer to release non-secreted proteins for pulldown. As a control, we also prepared lysates from WT bacteria that do not express tagged RARP-1. We then immunoprecipitated the lysates with a FLAG tag-specific antibody, performed an acid elution to release bound proteins, and analyzed the eluates by mass spectrometry to identify putative RARP-1 binding partners (Supplementary Figures 2A and B). Proteins that were present in the tagged lysate pulldown but absent from the untagged lysate pulldown were called as hits (Table 1 and Data Set 1).

**Table 1.**
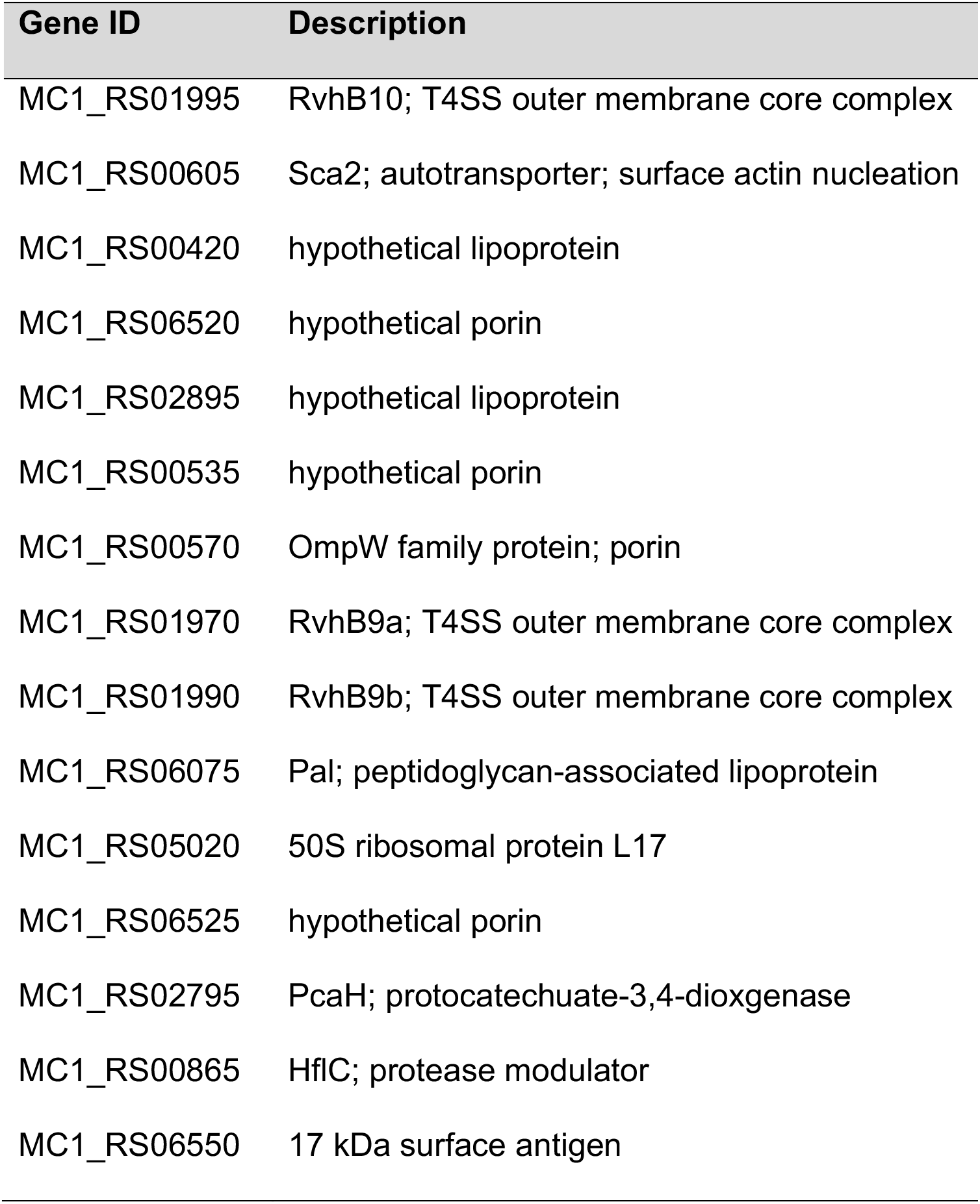
Co-immunoprecipitation of lysozyme-permeabilized bacteria reveals that RARP-1 interacts with other bacterial factors that access the periplasm.^a,b^. ^a^Putative RARP-1 binding partners are ordered by decreasing spectral count. ^b^MC1_RS05020 is the only hit not predicted to access the periplasm.

Of the hits identified, only Sca2 has been functionally characterized in *R. parkeri* (14). Although Sca2 promotes late-stage actin-based motility, the *rarp-1*::Tn mutant formed actin tails at frequencies comparable to WT (Figure 2A), indicating that the loss of RARP-1 does not dramatically impair Sca2 function. However, it is possible that RARP-1 functions in a more subtle way to influence Sca2 activity. To test this hypothesis, we used immunoblotting to assess Sca2 expression in the *rarp-1*::Tn mutant (Figure 6A). The abundance of full-length Sca2 and its processed products was comparable between the *rarp-1*::Tn mutant and the complement strain, suggesting that RARP-1 does not grossly impact Sca2 levels. Likewise, we observed similar patterns of Sca2 localization between strains (Figure 6B), suggesting that RARP-1 does not play a role in the polar positioning of Sca2. Taken together, these results suggest that RARP-1 does not regulate the activity of its putative binding partner Sca2.

**Figure 6.**
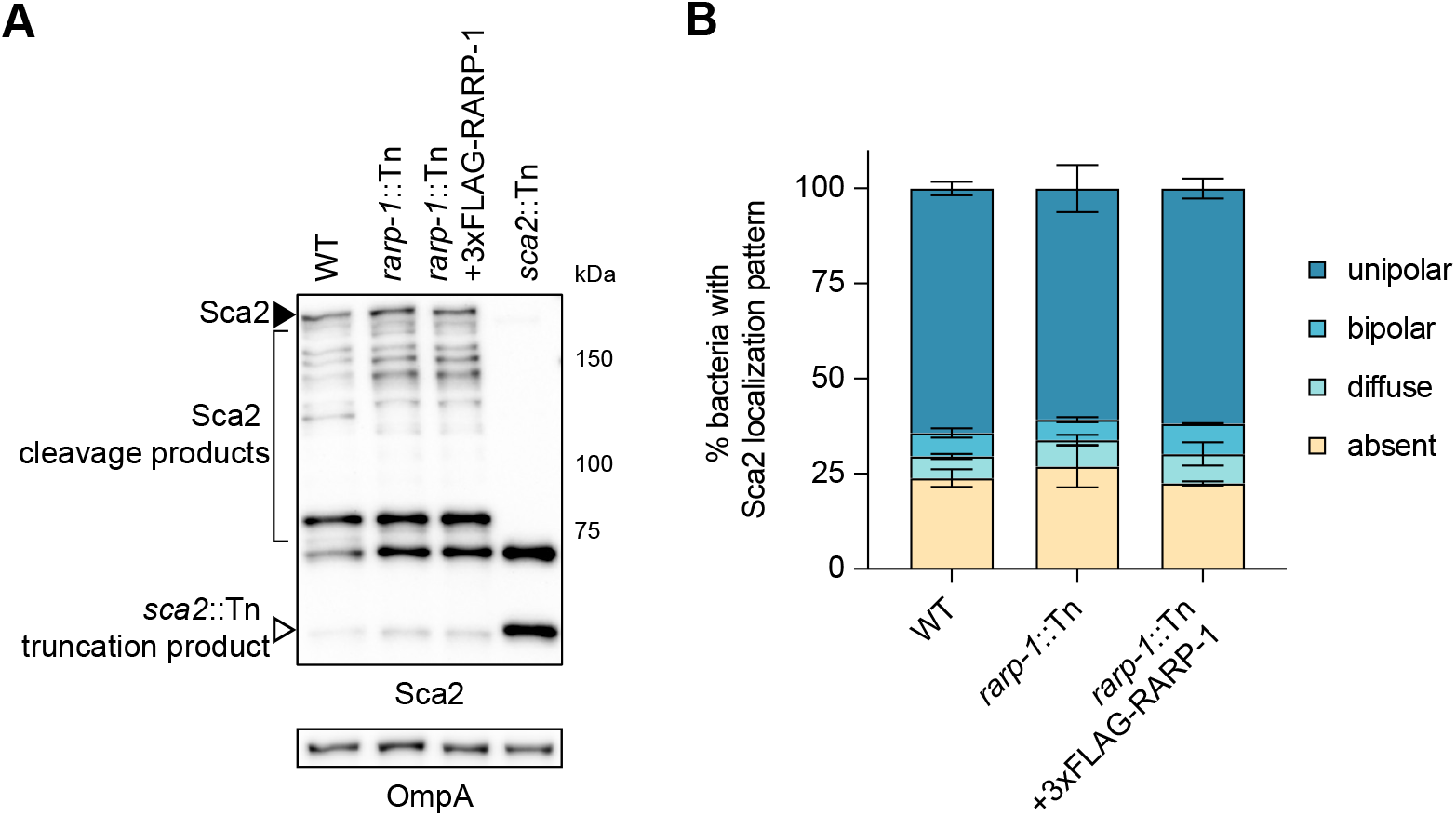
RARP-1 does not regulate the abundance or localization of Sca2. (A) Western blot for Sca2 from purified *R. parkeri* strains. Full-length Sca2 (arrowhead), Sca2 cleavage products (bracket), and the truncation product in the *sca2*::Tn mutant (open arrowhead) are indicated. (B) Percentage of bacteria with the indicated Sca2 localization pattern during infection of A549 cells. Percentages were determined from two independent experiments (≥ 350 bacteria were counted for each infection) and were used to calculate the mean ± SD and p-value (one-way ANOVA with post-hoc Dunnett’s test, n.s. relative to WT).

Additional hits identified in our analysis include the type IV secretion system outer membrane components RvhB9 and RvhB10 as well as several hypothetical lipoproteins and porins (Table 1). At this time, none of these proteins have been functionally characterized in *R. parkeri*. Consistent with RARP-1 localization to the periplasm, however, nearly all of these hits are predicted to reside in the periplasm or otherwise access and transit the periplasm *en route* to the bacterial surface. Thus, it remains possible that RARP-1 acts with one or more of these binding partners to support growth and host cell invasion.

## Discussion

After host cell invasion, obligate intracellular bacteria must scavenge host nutrients, proliferate, and avoid destruction by their hosts (2). Disruption of one or all of these activities will diminish intracellular bacterial loads and ultimately reduce pathogenicity. While many studies have revealed important regulators of invasion, nutrient acquisition, and bacterial growth for other species, little is known about the factors that support rickettsial physiology during infection, and only recently have we begun to uncover the protective strategies *Rickettsia* spp. employ to ward off host cell defenses (13, 16). Consequently, we sought to better understand the genetic determinants of rickettsial infection using our functional genetic approaches in *R. parkeri*. We found that RARP-1 likely resides in the periplasm where it interacts with proteins predicted or known to drive bacterial fitness or interactions with the host. Furthermore, our results suggest that RARP-1 supports the *R. parkeri* life cycle by promoting bacterial growth as well as efficient host cell invasion.

Loss of RARP-1 expression led to a transient invasion delay, suggesting that RARP-1 plays a role in host cell entry. Several studies have identified rickettsial surface proteins and candidate secreted effectors that facilitate invasion. For example, the outer membrane protein OmpA and OmpB respectively interact with □2β1 integrin and Ku70 at the host cell surface (5, 7), while the effectors RalF and Risk1 modulate host membrane phosphoinositides during entry (27, 28). Nevertheless, there is incomplete conservation across the *Rickettsia* genus for many of these proteins (27), and invasion is not abolished when the activity of any one protein is inhibited (13, 29); thus, it is likely that *Rickettsia* spp. use several redundant strategies to enter their hosts. Although RARP-1 itself is not exported from the bacterium, one or more of the RARP-1 interaction partners may contribute to efficient internalization as discussed above. Loss of RARP-1 expression would therefore have pleiotropic effects on infection by hindering both invasion and growth. Alternatively, it is possible that the *rarp-1*::Tn mutant invasion delay is the result of defective growth in the preceding infection cycles when the bacteria were harvested. Indeed, invasion competency of the intracellular bacterial pathogen *Brucella abortus* is linked to cell cycle progression (30). Perhaps rickettsial invasion efficiency relies on robust growth, without which the invasion program is impaired.

Loss of RARP-1 expression also reduced bacterial loads, persisting long after the initial invasion delay was overcome. This defect suggests that RARP-1 plays a role in bacterial growth through the regulation of bacterial physiology or avoidance of host defenses. Normally, *R. parkeri* shields itself from autophagy receptors by methylating outer membrane proteins such as OmpB (13). Loss of OmpB or the methyltransferases PKMT1 and PKMT2 promotes autophagy of *R. parkeri* and reduction of intracellular bacterial burdens (16). Since the *rarp-1*::Tn mutant did not display enhanced recruitment of the autophagy marker LC3, we concluded that the loss of RARP-1 does not render this mutant more susceptible to autophagy. Nevertheless, we cannot rule out that growth of the *rarp-1*::Tn mutant is restricted by other host defense strategies employed by the cell lines used in this study.

Prior work reported that RARP-1 was robustly secreted into the host cytoplasm by *R. typhi*, and experiments in *E. coli* suggested that RARP-1 relied on a non-canonical Sec- and TolC-dependent pathway for export (19). We were unable to detect secretion of endogenous or epitope-tagged RARP-1 into the host cytoplasm by *R. parkeri*, even though the tagged constructs functionally complemented the *rarp-1*::Tn mutant phenotype. Similarly, we were unable to detect phosphorylation of GSK-tagged RARP-1 in infected cell lysates as an orthogonal secretion assay. Notably, this lack of secretion was observed during infection of multiple host cell types and for both WT and *rarp-1*::Tn backgrounds. We also could not detect secretion of RARP-1 by *E. coli*, despite testing both *R. parkeri* and *R. typhi* homologs under the same conditions previously published (19). Nevertheless, it is formally possible that our use of a different *E. coli* K-12 strain (BW25513 rather than C600) prevented release of RARP-1 into the culture supernatant. Since *R. typhi* is a BSL-3 pathogen, we are not able to assess secretion of *R. typhi* RARP-1 by *R. parkeri*, and a loss-of-function *rarp-1* mutant does not exist in *R. typhi*. Altogether, our data suggest that RARP-1 is not secreted into the host cytoplasm by *R. parkeri*; instead, it is likely targeted to the periplasm by its Sec secretion signal where it stays to support bacterial growth and invasion.

RARP-1 is not predicted to possess enzymatic activity, but it does contain a large central intrinsically disordered region (IDR) and several C-terminal ankyrin repeats (ANKs). Although IDRs do not form ordered structures on their own, the structural plasticity of IDRs affords them diverse biological functions (31). For example, the IDRs of bacterial proteins facilitate chaperone recruitment, passage through narrow protein channels, and binding of multiple partners as part of a signaling hub (32–34). In *Caulobacter crescentus*, the IDR of PopZ serves as a scaffold for concentrating cell cycle regulators at the cytoplasmic cell poles (34). Given the localization pattern of RARP-1 and its interactions with various factors in the periplasm, it is possible that the IDR of RARP-1 performs a similar scaffolding role and concentrates binding partners at the *R. parkeri* periplasmic cell poles. Although our results suggest that the function and polar localization of the surface actin nucleator Sca2 is unaffected by the absence of RARP-1, additional studies will be necessary to assess the activity and localization of other RARP-1 binding partners in the *rarp-1*::Tn mutant.

ANKs are among the most common protein-protein interaction modules and ANK-containing proteins govern a variety of cellular processes (20). Many intracellular bacterial pathogens secrete ANK-containing effectors to target host cell functions, including protein trafficking, ubiquitination, and transcription (21, 22). Nevertheless, ANKs have also been shown to support the activity of bacterial proteins that are not secreted into the extracellular milieu. For example, AnkB localizes to the periplasm of *Pseudomonas aeruginosa* where it protects against oxidative stress (35), and Bd3460 of *Bdellovibrio bacteriovorus* complexes with endopeptidases in the periplasm to prevent degradation of its own cell wall (36). Although ANKs are best known for mediating protein-protein interactions, recent work has demonstrated that ANKs can also bind sugars and lipids (37, 38). Future mutational and biochemical analyses may reveal if the RARP-1 ANKs are necessary for interactions with its putative binding partners or if this domain also binds non-protein substrates in the periplasm to support RARP-1 activity.

Our data suggest that RARP-1 resides in the periplasm where it interacts with several classes of proteins to support growth and invasion. Since many of these binding partners have not been functionally characterized, we focused our attention on the interaction between RARP-1 and Sca2. Sca2 is required for late-stage actin-based motility in mammalian and tick cells (14, 39), and it is necessary for virulence in animal models of SFG rickettsial infection (40). Tn mutagenesis of *rarp-1* did not reduce actin tail frequency or Sca2 localization to the cell poles, suggesting that RARP-1 does not govern Sca2 function. Nevertheless, it is possible that Sca2 supports the localization or function of RARP-1 in the periplasm as it acts on other factors to regulate invasion and growth.

We also detected interactions between RARP-1 and components of the Rickettsiales *vir* homolog type IV secretion system (*rvh* T4SS). In the canonical *vir* T4SS of *Agrobacterium tumefaciens*, substrates are delivered from the bacterial cytoplasm into the host cell through a channel that spans the inner and outer membranes (41). VirB9 and VirB10, together with VirB7, form a core complex positioned in the periplasm and outer membrane (42). It is unknown to what extent the *rvh* subunits play similar roles as their *vir* counterparts, but it is possible that RARP-1 interacts with RvhB10 and both paralogs of RvhB9 in the periplasm to regulate T4SS assembly or export of effectors. At this time, few *rvh* T4SS effectors are known and none of them have been shown to modulate growth (23, 27, 28). Recent work has suggested that the putative *rvh* T4SS effector Risk1 promotes host cell invasion by *R. typhi* (28); whether Risk1 plays a similar role in *R. parkeri* or if its secretion is impacted in the *rarp-1*::Tn mutant is unknown. As new effectors are characterized, it will be important to determine if their secretion depends on the interaction between RARP-1 and the *rvh* T4SS. Alternatively, it is possible that the function or localization of RARP-1 is influenced by its interaction with RvhB9 and RvhB10. The T4SS of the intracellular bacterial pathogen *Legionella pneumophila* localizes to the poles (43); if the *rvh* T4SS behaves similarly, RARP-1 may be recruited to the periplasmic cell poles by its core complex binding partners.

Interestingly, many of the RARP-1 binding partners we identified include predicted porins (MC1_RS06520, MC1_RS00535, MC1_RS00570, and MC1_RS06525) and lipoproteins (MC1_RS00420 and MC1_RS02895) of unknown function, as well as the 17 kDa surface antigen. Porins are major components of the outer membrane and regulate the transport of hydrophilic compounds such as nutrients, toxins, and antibiotics (44). Homologs of MC1_RS00535 and MC1_RS00570 have been identified on the surface of the related SFG member *R. rickettsii* (45), but the substrates for these and other rickettsial porins have yet to be characterized. Lipoproteins are lipid-modified proteins that anchor to the membrane and support many aspects of bacterial physiology, including nutrient uptake, protein folding, signal transduction, and cell division (46). Based on remote homology predictions (via HHpred (47)), the hypothetical lipoproteins identified in this study appear to be unique to the *Rickettsia* genus and remain uncharacterized. Similarly, the 17 kDa surface antigen is unique to the genus and Tn mutagenesis of this gene reduces *R. parkeri* plaque size (18, 48), but its function is unknown. In future studies, it will be important to investigate how disruption of one or more of these factors contributes to the invasion and growth defects we observed for the *rarp-1*::Tn mutant.

The remaining RARP-1 interaction partners include homologs of proteins with known roles in bacterial physiology. For example, the peptidoglycan-associated lipoprotein Pal is concentrated at division septa through its interaction with the Tol machinery, which supports constriction of the outer membrane and remodeling of septal peptidoglycan during *E. coli* cell division (49). Although we did not observe any obvious morphological defects for the *rarp-1*::Tn mutant, the interaction between RARP-1 and Pal could influence rickettsial growth in a more subtle manner. PcaH, a subunit of protocatechuate-3,4-dioxygenase, was also identified as a RARP-1 binding partner. As part of the beta-ketoadipate pathway, this enzyme is involved in the conversion of aromatic compounds to TCA cycle intermediates (50). Homologs for all other enzymes in this pathway, however, are absent in the reduced genome of *R. parkeri* (via the KEGG pathway database (51)); thus, a role for PcaH and its interaction with RARP-1 during infection is unclear. Finally, we also detected an interaction between RARP-1 and HflC. HflC complexes with HflK in the periplasm to modulate the activity of the integral membrane protease FtsH (52). If RARP-1 provides an additional layer of regulation over FtsH through its interaction with HflC, it is possible that disruption of membrane protein quality control underlies the *rarp-1*::Tn mutant invasion and growth defects.

Our work uncovers an important role for RARP-1 in supporting the *R. parkeri* life cycle. Through its targeting to the periplasm, we propose that RARP-1 regulates invasion and growth by acting in concert with one or more of the factors revealed in our study. Further work is needed to characterize these interactions since many of the RARP-1 binding partners we identified have unknown functions in the *Rickettsia* genus. Expansion of the rickettsial toolkit could facilitate these efforts as well as help determine if there is temporal or spatial control of RARP-1 activity during the *R. parkeri* life cycle. Moreover, structure-function analyses of RARP-1 could provide valuable insight into its mechanism of action in particular and the function of ANK-and IDR-containing proteins in general. Homologs of RARP-1 are notably absent outside the genus, despite conservation of the protein across *Rickettsia* spp. (19). We therefore speculate that RARP-1 represents a core and unique adaptation to the demands of the host cell niche, and future studies may extend its relevance to infection of arthropod vectors. The success of *Rickettsia* spp. hinges on their ability to access and thrive within the complex environment of the host cytoplasm. Continued investigation into the factors that support these fundamental processes will not only improve our understanding of rickettsial biology, but will also highlight the diverse strategies underpinning obligate intracellular bacterial life.

## Materials and Methods

### Cell culture

A549 human lung epithelial and Vero monkey kidney epithelial cell lines were obtained from the University of California, Berkeley Cell Culture Facility (Berkeley, CA). A549 cells were maintained in DMEM (Gibco #11965118) containing 10% FBS. Vero cells were maintained in DMEM containing 5% FBS. A549 cells stably expressing cytoplasmic TagRFP-T (A549-TRT) were generated by retroviral transduction as previously described (15). Cell lines were confirmed to be mycoplasma-negative by MycoAlert PLUS Assay (Lonza #LT07-710) performed by the Koch Institute High Throughput Sciences Facility (Cambridge, MA).

### Plasmid construction

pRAM18dSGA-3xFLAG-RARP-1 was generated from pRAM18dSGA[MCS] (kindly provided by Dr. Ulrike Munderloh) and contains the 247 bp immediately upstream of the *tolC* start codon (MC1_RS01570), the first 23 aa (amino acids) of *R. parkeri* RARP-1 (MC1_RS01585) containing the Sec SS, a <u>H</u>VDYKDHDGDYKDHDIDYKDDDDKHV sequence (3xFLAG epitope tag underlined), the remaining 550 aa of RARP-1, and the *R. parkeri ompA* terminator (MC1_RS06480). pRL0079 is identical to pRAM18dSGA-3xFLAG-RARP-1 but contains <u>G</u>SGGEVH<u>TNQ</u>DPLDGGT (Ty1 epitope tag underlined) between residues 396 and 397.

pRL0284 was generated from pRAM18dSGA[MCS] and contains the *R. parkeri ompA* promoter, an N-terminal <u>MSGRPRTTSFAE</u>SGS sequence (GSK epitope tag underlined), TagBFP from pRAM18dRA-2xTagBFP (15), and the *ompA* terminator. pRL0285 is identical to pRL0284 but contains *R. parkeri* RARP-2 (MC1_RS04780) in place of TagBFP. Similarly, pRL0286 contains *R. parkeri* RARP-1 in place of TagBFP, but <u>G</u>SMSGRPRTTSFAESGS was inserted after the Sec SS (as in pRAM18dSGA-3xFLAG-RARP-1) instead of at the N-terminus.

pRL0287 was generated from pEXT20 (kindly provided by Dr. Michael Laub) and contains the *R. parkeri* RARP-1 insert with intervening 3xFLAG epitope tag from pRAM18dSGA-3xFLAG-RARP-1. pRL0288 is identical to pRL0287, except the 23 aa Sec SS of *R. typhi* RARP-1 (RT0218) and the remaining 563 aa of *R. typhi* RARP-1 were used. In contrast, pRL0289 contains the full 586 aa of *R. typhi* RARP-1 with a C-terminal <u>K</u>GEFEAYVEQKLISEEDLNSAV<u>D</u>HHHHHH sequence (Myc and 6xHis epitope tags underlined) as previously described (19). For pRL0290, a C-terminal <u>V</u>DHHHHHH sequence (6xHis epitope tag underlined) was added to *E. coli* YebF (NCBI b1847).

### Generation of *R. parkeri* strains

Parental *R. parkeri* str. Portsmouth (kindly provided by Dr. Chris Paddock) and all derived strains were propagated by infection and mechanical disruption of Vero cells grown in DMEM containing 2% FBS at 33 °C as previously described (15, 18). Bacteria were clonally isolated and expanded from plaques formed after overlaying infected Vero cell monolayers with agarose as previously described (18). When appropriate, bacteria were further purified by centrifugation through a 30% MD-76R gradient (Mallinckrodt Inc. #1317-07) as previously described (15). Bacterial stocks were stored as aliquots at −80 °C to minimize variability due to freeze-thaws. Titers were determined for bacterial stocks by plaque assay (15), and plaque sizes (Figure 1B) were measured with ImageJ after 5 d infection.

Bacteria were transformed with plasmids by small-scale electroporation as previously described (18), except infections were scaled down to a T25 cm^2^ flask and bacteria were electroporated with 1 μg dialyzed plasmid DNA. When appropriate, rifampicin (200 ng/mL) or spectinomycin (50 μg/mL) were included to select for transformants. The *rarp-1*::Tn and *sca2*::Tn mutants were generated as previously described (18), and the genomic locations of the Tn insertion sites were determined by semi-random nested PCR and Sanger sequencing. The expanded strains were verified by PCR amplification of the Tn insertion site using primers flanking the region. The *ompB^STOP^*::Tn mutant (referred to as *ompB*::Tn in this work; kindly provided by Dr. Matthew Welch) was generated as previously described (13).

### R. parkeri infections

For the infectious focus assays (Figures 2A and E and Supplementary Figures 1B and C), confluent A549 cells grown on 12 mm coverslips in 24-well plates were infected at an MOI of 0.005-0.025, centrifuged at 200 x g for 5 min at RT, and incubated at 33 °C for 1 h. Infected cells were washed three times with PBS before adding complete media with 10 μg/mL gentamicin. Infections progressed for 28 h at 33°C until fixation with 4% PFA in PBS for 10 min at RT.

To measure actin tail and protrusion frequencies (Figures 2B and C), confluent A549 cells grown on 12 mm coverslips in 24-well plates were infected at an MOI of 0.3-0.6, centrifuged at 200 x g for 5 min at RT, and incubated at 33 °C for 1 h. Infected cells were washed three times with PBS before adding complete media with 10 μg/mL gentamicin. Infections progressed for 28 h at 33°C until fixation with 4% PFA in PBS for 10 min at RT.

For the mixed-cell assays (Figure 2D), A549-TRT donor cells were plated in 96-well plates and unlabeled A549 recipient cells were plated in 6-well plates and grown to confluency. Donors were infected at an MOI of 9-10, centrifuged at 200 x g for 5 min at RT, and incubated at 33 °C for 1 h. Infected donors and uninfected recipients were washed with PBS, lifted with citric saline (135 mM KCl, 15 mM sodium citrate) at 37 °C to preserve cell surface receptors, recovered in complete media, washed twice with complete media to remove residual citric saline, and resuspended in complete media with 10 μg/mL gentamicin (6 x 10^5^ cells/mL donors and 8 x 10^5^ cells/mL recipients). Cells were then mixed at a 1:125 ratio (5.3 μL donors and 500 μL recipients) and plated on 12 mm coverslips in 24-well plates. Infections progressed for 31 h at 33 °C until fixation with 4% PFA in PBS for 1 h at RT.

To measure growth (Figure 2F), confluent Vero cells grown in 24-well plates were infected in triplicate at an MOI of 0.025, centrifuged at 200 x g for 5 min at RT, and incubated at 33 °C for 1 h. Infected cells were washed three times with serum-free DMEM before adding complete media and allowing infections to progress at 33 °C. To harvest samples at the indicated time point, infected cells were scraped into the media and centrifuged at 20,000 x g for 5 min. The resulting pellets were resuspended in 600 μL Nuclei Lysis Solution (Promega #A7941), boiled for 10 min to release genomic DNA, and processed with a Wizard Genomic DNA Purification Kit (Promega #A1125) according to manufacturer instructions. After air-drying, the DNA pellets were resuspended in 100 μL H_2_O, incubated at 65 °C for 1 h, and allowed to completely rehydrate overnight at RT. For qPCR, runs were carried out on a LightCycler 480 (Roche) at the MIT BioMicro Center (Cambridge, MA). Primers to the *R. parkeri* 17 kDa surface antigen gene (MC1_RS06550; 5’-TTCGGTAAGGGCAAAGGACA-3’ and 5’-GCACCGATTTGTCCACCAAG-3’) and to *Chlorocebus sabaeus GAPDH* (5’-AATGGGACTGAAGCTCCTGC-3’ and 5’-ATCACCACCCCTCTACCTCC-3’) were used to determine bacterial and host genome equivalents, respectively, relative to a standard curve prepared from a pooled mixture of the 96 h time point WT infection samples. Results from each biological replicate were normalized to the 1 h time point and fold-change was calculated.

To evaluate LC3 recruitment (Figure 3A), confluent A549 cells grown on 12 mm coverslips in 24-well plates were infected at an MOI of 1.8-3.6, centrifuged at 200 x g for 5 min at RT, and incubated at 33 °C for 2 h until fixation with 4% PFA in PBS for 10 min.

To measure invasion efficiency (Figure 3B), confluent A549 cells grown on 12 mm coverslips in 24-well plates were placed on ice and the media was replaced with 500 μL ice-cold complete media. The cells were then infected at an MOI of 0.7-1.2, centrifuged at 200 x g for 5 min at 4 °C, 500 μL 37 °C complete media was added, and the plates were immediately moved to 37 °C until fixation with 4% PFA in PBS for 10 min.

To evaluate secretion of RARP-1 (Figure 4A and Supplementary Figures 1A and D), confluent A549 cells grown in 24-well plates were infected at an MOI of 0.5-1.0, centrifuged at 200 x g for 5 min at RT, and incubated at 33 °C until the indicated harvest time point (Figure 4A) or for 48 h (Supplementary Figures 1A and D).

To evaluate secretion of GSK-tagged constructs (Figure 4B), confluent Vero cells grown in 24-well plates were infected with the indicated strains, centrifuged at 200 x g for 5 min at RT, and incubated at 33 °C with spectinomycin for 72 h (when infected cells were approximately 90% rounded) before harvesting.

To evaluate the localization of epitope-tagged RARP-1 (Figures 5A-C), confluent A549 cells grown in 24-well plates were infected at an MOI of 0.3-0.6, centrifuged at 200 x g for 5 min at RT, and incubated at 33 °C for 27 h until fixation with 4% PFA in PBS for 1 h.

To evaluate the localization of Sca2 (Figure 6B), confluent A549 cells grown in 24-well plates were infected at an MOI of 0.3-0.6, centrifuged at 200 x g for 5 min at RT, and incubated at 33 °C for 28 h until fixation with 4% PFA in PBS for 10 min.

### *E. coli* secretion assays

*E. coli* K-12 BW25113 (WT) and JW5503-1 (Δ*tolC*) from the Keio Knockout Collection (53) were obtained from Horizon Discovery. SDS sensitivity and the KanR cassette insertion site were confirmed for the Δ*tolC* strain. Secretion assay samples were collected and processed as previously described (19). Bacterial pellets and precipitated proteins were boiled in loading buffer (50 mM Tris-HCl pH 6.8, 2% SDS, 10% glycerol, 0.1% bromophenol blue, 5% β-mercaptoethanol).

### RARP-1 antibody production

The RARP-1 peptide antigen (SNEMHEAQVASNEHND, corresponding to residues 159-174) was selected and synthesized by New England Peptide (Gardner, MA). The peptide antigen was conjugated to KLH and used for immunization by Pocono Rabbit Farm and Laboratory (Canadensis, PA) according to their 70 day rabbit polyclonal antibody protocol.

### Immunoblotting

To assess RARP-1 and Sca2 expression (Figures 1C, 1D, and 6A), purified bacteria were boiled in loading buffer and analyzed by western blot using rabbit RARP-1 peptide antisera, rabbit anti-FLAG (Cell Signaling Technology #2368), rabbit anti-Sca2 (kindly provided by Dr. Matthew Welch), and mouse anti-OmpA 13-3 (kindly provided by Dr. Ted Hackstadt). For Figures 1C and 6A, the parental WT *R. parkeri* strain lacking pRAM18dRGA+OmpApr-GFPuv was used. In Figures 1C and D, the apparent MW of RARP-1 is greater than its predicted MW (60 kDa). This aberrant migration by SDS-PAGE is typical of proteins with IDRs (54).

To evaluate secretion of RARP-1 (Figure 4A and Supplementary Figures 1A and D), infected cells were washed three times with PBS, lifted with trypsin-EDTA, and centrifuged at 2,400 x g for 5 min at RT. The resulting pellets were resuspended in selective lysis buffer (50 mM HEPES pH 7.9, 150 mM NaCl, 1 mM EDTA, 10% glycerol, 1% IGEPAL) containing protease inhibitors (Sigma-Aldrich #P1860), incubated on ice for 15 min, and centrifuged at 11,300 x g for 10 min at 4 °C. The resulting pellets were washed with PBS and boiled in loading buffer. The resulting supernatants were passed through a 0.22 μm cellulose acetate filter (Thermo Scientific #F2517-1) by centrifugation at 6,700 x g for 10 min at 4 °C, combined with loading buffer (to a final volume equal to the final pellet volume), and boiled. Lysates were analyzed by western blot using rabbit anti-FLAG, rabbit anti-Sca4 (15), mouse anti-Ty1 (kindly provided by Dr. Sebastian Lourido), rabbit RARP-1 peptide antisera, and mouse anti-RpoA (BioLegend #663104).

To evaluate secretion of GSK-tagged constructs (Figure 4B), infected cells were washed with ice-cold serum-free DMEM, directly lysed in loading buffer, and boiled. Lysates were analyzed by western blot using rabbit anti-GSK-3β-Tag (Cell Signaling Technology #9325) and rabbit anti-phospho-GSK-3β (Cell Signaling Technology #9336).

For the *E. coli* secretion assays (Figures 4C and D), bacterial pellet lysates (equivalent to 0.025 OD_600_-mL of cultured cells) and precipitated culture supernatants (equivalent to 2 mL of culture supernatant prior to precipitation) were analyzed by western blot using rabbit anti-FLAG and HRP-conjugated mouse anti-His (ABclonal #AE028).

For the co-immunoprecipitation assays (Supplementary Figures 2A and B), samples were analyzed by western blot using rabbit anti-Sca4 and rabbit anti-FLAG.

### Immunofluorescence microscopy

All micrographs were acquired on an Olympus IXplore Spin microscope system, and image analysis was performed with ImageJ unless otherwise stated.

For the infectious focus assays (Figures 2A and E and Supplementary Figures 1B and C), fixed samples were incubated with 0.1 M glycine in PBS for 10 min at RT to quench residual PFA. Samples were then washed three times with PBS, permeabilized with 0.5% Triton X-100 in PBS for 5 min at RT, and washed another three times with PBS. Samples were then incubated with blocking buffer (2% BSA in PBS) for 30 min at RT. Primary and secondary antibodies were diluted in blocking buffer and incubated for 1 h each at RT with three 5 min PBS washes after each incubation step. The following antibodies and stains were used: mouse anti-β-catenin (Cell Signaling Technology #2677) to detect host membrane, rabbit anti-*Rickettsia* I7205 (kindly provided by Dr. Ted Hackstadt), goat anti-mouse conjugated to Alexa Fluor 568 (Invitrogen #A-11004), goat anti-rabbit conjugated to Alexa Fluor 488 (Invitrogen #A-11008), and Hoechst (Invitrogen #H3570) to detect host nuclei. Coverslips were mounted using ProLong Gold Antifade Mountant (Invitrogen #P36934). Images were acquired using a 60X UPlanSApo (1.30 NA) objective. For each strain, 20-35 foci were imaged and the number of infected cells and bacteria per focus was calculated.

To measure actin tail and protrusion frequencies (Figures 2B and C), fixed samples were processed as above, except phalloidin conjugated to Alexa Fluor 647 (Invitrogen #A22287) was included to detect actin. For each strain, ≥ 380 bacteria were imaged using a 100X UPlanSApo (1.35 NA) objective and the percentage of bacteria with tails (> 1 bacterial length) and the percentage of bacteria within protrusions were calculated.

For the mixed-cell assays (Figure 2D), fixed samples were processed as above, except the following antibodies and stains were used: mouse anti-*Rickettsia* 14-13 (kindly provided by Dr. Ted Hackstadt), goat anti-mouse conjugated to Alexa Fluor 647 (Invitrogen #A-21235), and phalloidin-iFluor 405 Reagent (Abcam #ab176752). For each strain, 20 foci were imaged using a 60X objective and the percentage of bacteria per focus that had spread to recipient cells was calculated.

To evaluate LC3 recruitment (Figure 3A), fixed samples were processed as above, except cells were permeabilized with 100% methanol for 5 min at RT instead of Triton X-100 and the following antibodies and stains were used: rabbit anti-LC3B (ABclonal #A7198), mouse anti-*Rickettsia* 14-13, goat anti-rabbit conjugated to Alexa Fluor 568, goat anti-mouse conjugated to Alexa Fluor 488, and Hoechst. Representative images were acquired using a 100X objective.

To measure invasion efficiency (Figure 3B), fixed samples were incubated with 0.1 M glycine in PBS for 10 min at RT to quench residual PFA. Samples were then washed three times with PBS and incubated with blocking buffer for 30 min at RT. To stain external bacteria, primary and secondary antibodies were diluted in blocking buffer and incubated for 30 min each at RT with three 5 min PBS washes after each incubation step. The following antibodies and stains were used: mouse anti-*Rickettsia* 14-13 and goat anti-mouse conjugated to Alexa Fluor 647. The samples were then fixed with 4% PFA in PBS for 5 min at RT, washed three times with PBS, and quenched with 0.1 M glycine in PBS for 10 min at RT. Samples were then washed three times with PBS, permeabilized with 0.5% Triton X-100 in PBS for 5 min at RT, and washed another three times with PBS. To stain both external and internal bacteria, primary and secondary antibodies were diluted in blocking buffer and incubated for 30 min each at RT with three 5 min PBS washes after each incubation step. The following antibodies and stains were used: mouse anti-*Rickettsia* 14-13 and goat anti-mouse conjugated to Alexa Fluor 488. For each strain, 20 fields of view each containing ≥ 45 bacteria were imaged using a 60X objective. To facilitate analysis, internal and external bacteria were quantified using ilastik (55); the pixel classifier was trained to distinguish bacteria from background, and then the object classifier was trained to distinguish between internal (single-stained) and external (double-stained) bacteria.

To evaluate the localization of epitope-tagged RARP-1 (Figures 5A-C), fixed samples were incubated with 0.1 M glycine in PBS for 10 min at RT to quench residual PFA. Samples were then washed three times with PBS, permeabilized with 0.5% Triton X-100 in PBS for 5 min at RT, and washed another three times with PBS. Samples were then incubated with goat serum blocking buffer (2% BSA and 10% normal goat serum in PBS) for 30 min at RT. To stain host cell contents and bacterial surface proteins, primary and secondary antibodies were diluted in goat serum blocking buffer and incubated for 3 h at 37 °C and 1 h at RT, respectively, with three 5 min PBS washes after each incubation step. For Figure 5A, rabbit anti-FLAG, mouse anti-*Rickettsia* 14-13, goat anti-rabbit conjugated to Alexa Fluor 647 (Invitrogen #A-21245), and goat anti-mouse conjugated to Alexa Fluor 488 were used, and coverslips were mounted after washing. For Figure 5B, only mouse anti-*Rickettsia* 14-13 and goat anti-mouse conjugated to Alexa Fluor 488 were used in the first round of staining, and coverslips were instead fixed with 4% PFA in PBS for 5 min at RT after washing. These samples were then incubated with 0.1 M glycine in PBS for 10 min at RT to quench residual PFA and washed three times with PBS. To expose proteins inside the bacteria for staining, these samples were incubated with lysozyme reaction buffer (0.8X PBS, 50 mM glucose, 5 mM EDTA, 0.1% Triton X-100, 5 mg/mL lysozyme (Sigma #L6876)) for 20 min at 37 °C and then washed three times with PBS. Rabbit anti-FLAG and goat anti-rabbit conjugated to Alexa Fluor 647 were diluted in goat serum blocking buffer and incubated for 3 h at 37 °C and 1 h at RT, respectively, with three 5 min PBS washes after each incubation step. Coverslips were mounted after the second round of staining. For Figure 5C, the same procedure was used as in Figure 5B, except goat anti-mouse conjugated to Alexa Fluor 488 was replaced with Alexa Fluor 405 (Invitrogen #A-31553) to permit imaging of bacterial GFP. Representative images were acquired using a 100X objective. Images in Figure 5C were deconvolved by performing five iterations of the cellSens (Olympus) advanced maximum likelihood estimation algorithm, and a 0.26 μm width pole-to-pole linescan was performed with ImageJ.

To evaluate the localization of Sca2 (Figure 6B), fixed samples were processed as above, except the following antibodies and stains were used: rabbit anti-Sca2, mouse anti-*Rickettsia* 14-13, goat anti-rabbit conjugated to Alexa Fluor 568 (Invitrogen #A-11011), goat anti-mouse conjugated to Alexa Fluor 488 (Invitrogen #A-11001), phalloidin conjugated to Alexa Fluor 647, and Hoechst. For each strain, ≥ 350 bacteria were imaged using a 100X objective and the Sca2 localization pattern was determined (following the classification scheme from (14)).

### Co-immunoprecipitation assays

Two replicate samples each of WT and *rarp-1*::Tn + 3xFLAG-Ty1-RARP-1 bacteria were processed in parallel for FLAG co-immunoprecipitation. For each sample, bacteria purified from a fully infected T175 cm^2^ flask were centrifuged at 16,200 x g for 2 min at RT, resuspended in 1 mL immunoprecipitation lysis buffer (50 mM Tris-HCl pH 7.4, 150 mM NaCl, 1 mM EDTA, 1% Triton X-100) containing 50 U/μL Ready-Lyse Lysozyme (Lucigen #R1804M) and protease inhibitors, incubated for 25 min at RT, and centrifuged at 11,300 x g for 15 min at 4 °C. The resulting supernatants were pre-cleared twice by incubation with 28 μL 50% mouse IgG agarose slurry (Sigma #A0919) for 30 min at 4°C. The pre-cleared input lysates were then incubated with 28 μL 50% anti-FLAG M2 magnetic bead slurry (Sigma #M8823) overnight at 4 °C. The bound complexes were washed four times with 500 μL ice-cold immunoprecipitation wash buffer (50 mM Tris-HCl pH 7.4, 150 mM NaCl) containing protease inhibitors, eluted by incubation with 65.2 μL 0.1 M glycine (pH 2.8) for 20 min at RT, and neutralized with 9.8 μL 1 M Tris-HCl (pH 8.5). The neutralized eluates were then combined with loading buffer and submitted to the Whitehead Institute Proteomics Core Facility (Cambridge, MA) for sample workup and mass spectrometry analysis. Equivalent bacterial input was confirmed by immunoblotting for Sca4 (Supplementary Figure 2A).

### Mass spectrometry

Samples were run 1 cm into an SDS-PAGE gel, excised, and then reduced, alkylated, and digested with trypsin overnight at 37 °C. The resulting peptides were extracted, concentrated, and injected onto a nanoACQUITY UPLC (Waters) equipped with a self-packed Aeris 3.6 μm C18 analytical column (20 cm x 75 μm; Phenomenex). Peptides were eluted using standard reverse-phase gradients. The effluent from the column was analyzed using an Orbitrap Elite mass spectrometer (nanospray configuration; Thermo Scientific) operated in a data-dependent manner. Peptides were identified using SEQUEST (Thermo Scientific) and the results were compiled in Scaffold (Proteome Software). RefSeq entries for *R. parkeri* str. Portsmouth (taxonomy ID 1105108) and *Homo sapiens* (taxonomy ID 9606) were downloaded from NCBI and concatenated with a database of common contaminants. Peptide identifications were accepted at a threshold of 95%. Protein identifications were accepted with a threshold of 99% and two unique peptides. Rickettsial proteins that were present in both replicates of the tagged (*rarp-1*::Tn + 3xFLAG-Ty1-RARP-1) lysate pulldown but absent from both replicates of the untagged (WT) lysate pulldown were called as hits.

### Statistical analyses

Statistical analysis was performed using Prism 9 (GraphPad Software). Graphical representations, statistical parameters, and significance are reported in the figure legends. Data were considered to be statistically significant when p < 0.05, as determined by an unpaired Student’s *t* test or one-way ANOVA with post-hoc Dunnett’s test.

### Data Availability

Mass spectral data and the protein sequence database used for searches have been deposited in the public proteomics repository MassIVE (https://massive.ucsd.edu, MSV000088867).

## Acknowledgements

We are grateful to Michael Laub, Jon McGinn, and Brandon Sit for critical review of the manuscript. We thank Ted Hackstadt, Michael Laub, Sebastian Lourido, Ulrike Munderloh, Chris Paddock, and Matthew Welch for reagents and Sebastian Lourido, Elizabeth Boydston, Natasha Kafai, and Adam Nock for technical help. We also thank Whitehead Institute Proteomics Core Facility members Eric Spooner and Edward Dudek for experimental support. This work was supported by NIH/NIGMS T32GM007287 and T32GM136540 (A.G.S., R.E.H.), NIH/NIGMS R00GM115765 (R.L.L.), and NIH/NIAID R01AI155489 (R.L.L.).

## Supplemental Material

**Supplementary Figure 1. Tagged RARP-1 constructs and endogenous RARP-1 are not secreted.** (A) *R. parkeri* RARP-1 with insertion sites for 3xFLAG and Ty1 epitope tags indicated (arrowheads). Western blots for FLAG (top) and Ty1 (middle) after infection of A549 cells with *rarp-1*::Tn + 3xFLAG-RARP-1 (single-tagged) or *rarp-1*::Tn + 3xFLAG-Ty1-RARP-1 (dual-tagged) bacteria. (B) Infected cells per focus during infection of A549 cells. (C) Bacteria per focus during infection of A549 cells. In (B) and (C), the means from three independent experiments (squares) are superimposed over the raw data (circles) and were used to calculate the mean ± SD and p-value (one-way ANOVA with post-hoc Dunnett’s test, **p < 0.01 relative to WT). (D) Western blots for RARP-1 (top) and Sca4 (middle) after infection of A549 cells with WT or *rarp-1*::Tn bacteria. Note the specific RARP-1 band in the pellet sample for WT bacteria only, in contrast to the identical non-specific bands in the supernatant samples for WT and *rarp-1*::Tn bacteria. In (A) and (D), infected host cells were selectively lysed after 48 h to separate supernatants (S) containing the infected host cytoplasm from pellets (P) containing intact bacteria. RpoA (bottom) served as a control for bacterial lysis or contamination of the infected cytoplasmic fraction.

**Supplementary Figure 2. Inputs and eluates from co-immunoprecipitation of lysozyme-permeabilized bacteria.** (A) Western blot for Sca4 (loading control) in input lysates. (B) Western blot for FLAG in input lysates and FLAG immunoprecipitation eluates. In (A) and (B), bacteria expressing tagged (+) or untagged (–) RARP-1 were purified and then permeabilized by lysozyme prior to immunoprecipitation. Two replicate samples were harvested from each strain.

**Data Set 1. Full co-immunoprecipitation / mass spectrometry results.**

**Supplementary Table 1. Strains and plasmids used in this study.**

