## Supplementary material for "The ankyrin repeat protein RARP-1 is a periplasmic factor that supports *Rickettsia parkeri* growth and host cell invasion": Figure S1

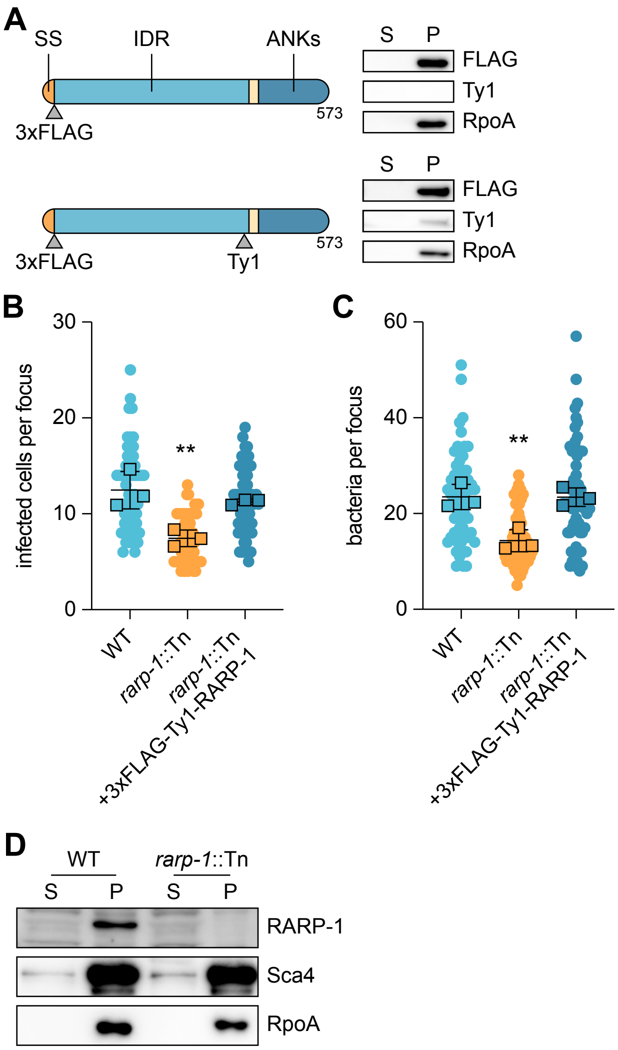


**Supplementary Figure 1. Tagged RARP-1 constructs and endogenous RARP-1 are not secreted.** (A) *R. parkeri* RARP-1 with insertion sites for 3xFLAG and Ty1 epitope tags indicated (arrowheads). Western blots for FLAG (top) and Ty1 (middle) after infection of A549 cells with *rarp-1*::Tn + 3xFLAG-RARP-1 (single-tagged) or *rarp-1*::Tn + 3xFLAG-Ty1-RARP-1 (dual-tagged) bacteria. (B) Infected cells per focus during infection of A549 cells. (C) Bacteria per focus during infection of A549 cells. In (B) and (C), the means from three independent experiments (squares) are superimposed over the raw data (circles) and were used to calculate the mean ± SD and p-value (one-way ANOVA with post-hoc Dunnett's test, $^{**}p < 0.01$ relative to WT). (D) Western blots for RARP-1 (top) and Sca4 (middle) after infection of A549 cells with WT or *rarp-1*::Tn bacteria. Note the specific RARP-1 band in the pellet sample for WT bacteria only, in contrast to the identical non-specific bands in the supernatant samples for WT and *rarp-1*::Tn bacteria. In (A) and (D), infected host cells were selectively lysed after 48 h to separate supernatants (S) containing the infected host cytoplasm from pellets (P) containing intact bacteria. RpoA (bottom) served as a control for bacterial lysis or contamination of the infected cytoplasmic fraction.
