## Supplementary material for "The ankyrin repeat protein RARP-1 is a periplasmic factor that supports *Rickettsia parkeri* growth and host cell invasion": Figure S2

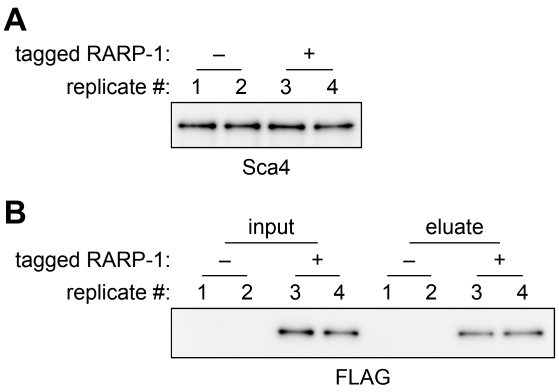


**Supplementary Figure 2. Inputs and eluates from co-immunoprecipitation of lysozyme-permeabilized bacteria.** (A) Western blot for Sca4 (loading control) in input lysates. (B) Western blot for FLAG in input lysates and FLAG immunoprecipitation eluates. In (A) and (B), bacteria expressing tagged (+) or untagged (–) RARP-1 were purified and then permeabilized by lysozyme prior to immunoprecipitation. Two replicate samples were harvested from each strain.
