## Supplementary material for "The ankyrin repeat protein RARP-1 is a periplasmic factor that supports *Rickettsia parkeri* growth and host cell invasion": Table S1

| Strain or plasmid | Genotype or feature | Reference or source |
|---|---|---|
| ***R. parkeri* strains** | | |
| *R. parkeri* str. Portsmouth | Parental *R. parkeri* strain | Chris Paddock |
| WT | pRAM18dSGA+OmpApr-GFPuv | (15) |
| *rarp-1*::Tn | *rarp-1*::Tn | (18) |
| *sca2*::Tn | *sca2*::Tn + pRAM18dSGA+OmpApr-GFPuv | (18) |
| *ompB*$^{STOP}$::Tn | *ompB*$^{STOP}$::Tn | (13) |
| *rarp-1*::Tn + 3xFLAG-RARP-1 | *rarp-1*::Tn + pRAM18dSGA-3xFLAG-RARP-1 | This study |
| *rarp-1*::Tn + 3xFLAG-Ty1-RARP-1 | *rarp-1*::Tn + pRL0079 | This study |
| GSK-BFP | pRL0284 | This study |
| GSK-RARP-2 | pRL0285 | This study |
| GSK-RARP-1 | pRL0286 | This study |
| ***E. coli* strains** | | |
| WT | Keio Knockout Collection parental K12 strain (BW25113) | (53); Horizon Discovery |
| Δ*tolC* | Δ*tolC*::Kan (JW5503-1) | (53); Horizon Discovery |
| WT + 3xFLAG-RARP-1$_{Rp}$ | WT + pRL0287 | This study |

| | | |
|---|---|---|
| WT + 3xFLAG-RARP-$1_{Rt}$ | WT + pRL0288 | This study |
| Δ*tolC* + 3xFLAG-RARP-$1_{Rp}$ | Δ*tolC* + pRL0287 | This study |
| Δ*tolC* + 3xFLAG-RARP-$1_{Rt}$ | Δ*tolC* + pRL0288 | This study |
| WT + Myc-6xHis-RARP-$1_{Rt}$ | WT + pRL0290 | This study |
| WT + 6xHis-YebF | WT + pRL0291 | This study |
| **Plasmids** | | |
| pRAM18dSGA[MCS] | *Rickettsia* shuttle vector | Ulrike Munderloh |
| pRAM18dSGA+OmpApr-GFPuv | GFPuv | (15) |
| pRAM18dSGA-3xFLAG-RARP-1 | 3xFLAG-tagged *R. parkeri* RARP-1 | This study |
| pRL0079 | 3xFLAG- and Ty1-tagged *R. parkeri* RARP-1 | This study |
| pRL0284 | GSK-tagged TagBFP | This study |
| pRL0285 | GSK-tagged *R. parkeri* RARP-2 | This study |
| pRL0286 | GSK-tagged *R. parkeri* RARP-1 | This study |
| pEXT20 | IPTG-inducible *E. coli* expression vector | This study |
| pRL0287 | 3xFLAG-tagged *R. parkeri* RARP-1 | This study |

| | | |
|---|---|---|
| pRL0288 | 3xFLAG-tagged *R. typhi* RARP-1 | This study |
| pRL0290 | Myc-6xHis-tagged *R. typhi* RARP-1 | This study |
| pRL0291 | 6xHis-tagged *E. coli* YebF | This study |

**Supplementary Table 1. Strains and plasmids used in this study.**
